# Robust Neutralization of SARS-CoV-2 Variants Including JN.1 and BA.2.87.1 by Trivalent XBB Vaccine-Induced Antibodies

**DOI:** 10.1101/2024.02.16.580615

**Authors:** Xun Wang, Shujun Jiang, Wentai Ma, Chen Li, Changyi Liu, Faren Xie, Jinsheng Zhu, Yan Zhan, Shibo Jiang, Mingkun Li, Yanliang Zhang, Pengfei Wang

## Abstract

Newly emerged SARS-CoV-2 variants like JN.1, and more recently, the hypermutated BA.2.87.1, have raised global concern. We recruited two groups of participants who had BA.5/BF.7 breakthrough infection post three doses of inactivated vaccines: one group experienced subsequent XBB reinfection, while the other received the XBB-containing trivalent WSK-V102C vaccine. Our comparative analysis of their serum neutralization activities revealed that the WSK-V102C vaccine induced stronger antibody responses against a wide range of variants, notably including JN.1 and the highly escaped BA.2.87.1. Furthermore, our investigation into specific mutations revealed that fragment deletions in NTD significantly contribute to the immune evasion of the BA.2.87.1 variant. Our findings emphasize the necessity for ongoing vaccine development and adaptation to address the dynamic nature of SARS-CoV-2 variants.

## INTRODUCTION

The rapid and widespread transmission of SARS-CoV-2 has led to the emergence of multiple variants of concern (VOCs), most notably Omicron variant, which continues to evolve and diversify into a range of sub-lineages. Our previous research has shown that these Omicron sub-lineages, spanning BA.1 to BA.2.86, have been evolving to exhibit increased neutralization escape capabilities^1-5^. Since November 2023, the JN.1 variant, stemming from BA.2.86’s antigenic diversity and acquiring an additional mutation (L455S) in RBD, has rapidly emerged as the dominant strain (Fig. 1a). Additionally, a newly identified, highly mutated SARS-CoV-2 variant, BA.2.87.1, was detected in South Africa between September and November 2023, which has recently been classified as a variant under monitoring (VUM) and has sparked global concern. While it originates from the ancestral BA.2 lineage, BA.2.87.1 is genetically distinct from the currently circulating Omicron lineages (Fig. 1b). It exhibits over 100 mutations, with more than 30 in the spike protein (Supplementary Table 1), including notable changes in the receptor binding domain (RBD) like K417T, K444N, V445G, and L452M, which are crucial for antibody recognition. Intriguingly, this variant has 7 fragment deletions, including 3 in the spike protein, with 2 of which encompasses over 10 crucial amino acids (Del 15-26 and Del 136-146) in the N-terminal domain (NTD) (Fig. 1c, d). This evolutionary strategy, which involves sacrificing parts of the virus to evade the immune system, makes these deletions potentially more significant than nonsynonymous mutations. Given our existing immunity from vaccinations and past infection, the effectiveness of this immunity against BA.2.87.1 remains to be determined.

**Fig. 1.**
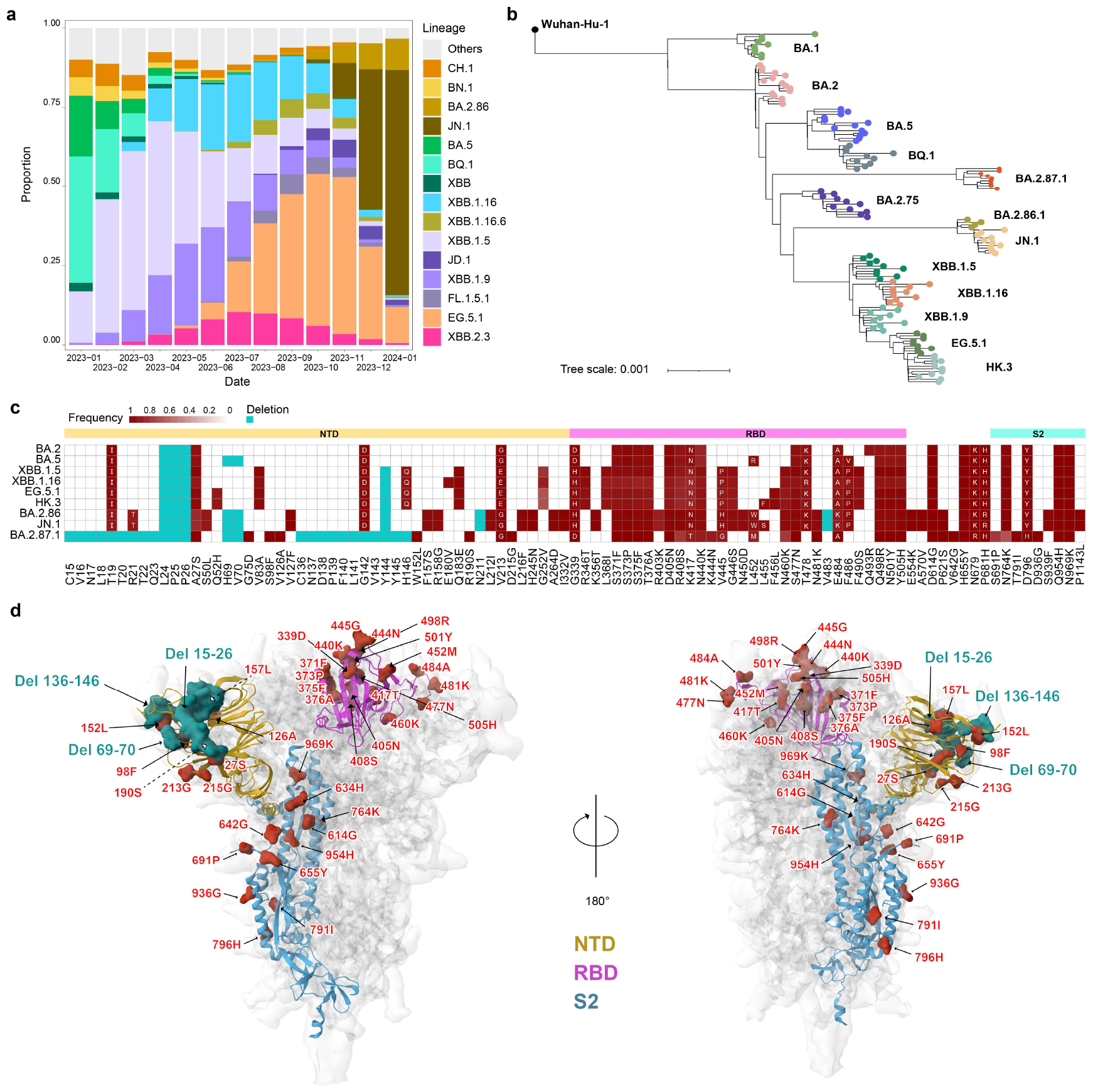
Characteristics of the recently emerging BA.2.87.1 variant. **a** The relative frequencies of SARS-CoV-2 lineages over time. Data were obtained from the CNCB RCoV19 database (ngdc.cncb.ac.cn/ncov/). **b** Phylogenetic tree of BA.2.87.1 and other Omicron sub-lineages. Sequences were sub-sampled from the Nextstrain GISAID global dataset of the last 6 months (updated on 2024 Feb 13). **c** The mutation frequency heatmap of BA.2.87.1 and other related lineages. Only mutations with a frequency higher than 0.75 are shown. Deletions are shown in cyan. Mutation frequency data was retrieved from the outbreak.info website. **d** The mutations of BA.2.87.1 on the SARS-CoV-2 spike glycoprotein trimer (PDB: 7WGV). Deletions are represented by the cyan molecular surface, while SNPs are represented by the red molecular surface. The NTD, RBD, and S2 regions were marked in yellow, pink, and blue, respectively.

The swift rise of antigenically diverse SARS-CoV-2 variants and decreasing vaccine efficacy against infection have necessitated updates in COVID-19 vaccine formulations. In September 2023, the monovalent XBB.1.5 mRNA vaccine received approval from the United States Food and Drug Administration (FDA). Concurrently, China approved several XBB-adapted vaccines, both multivalent and monovalent, including SYS6006 (CSPC Pharmaceutical Group), RQ3033 (Watson Biotech), SCTV01E-2 (SinocellTech), and WSK-V102C (WestVac Biopharma). The WestVac BioPharma’s Coviccine® Trivalent XBB Vaccine (WSK-V102C) incorporates the RBDs of the XBB.1.5, BA.5, and Delta variants, fused with the spike protein’s heptad repeat (HR) domain and self-assembling into stable trimeric protein particles. The vaccine is further enhanced with a squalene-based oil-in-water emulsion adjuvant, added post-purification and mixing for increased efficacy^6^.

From December 2022 to January 2023, over 80% of China’s population experienced BA.5/BF.7 infections despite receiving three doses of inactivated vaccines. Subsequently, from May to July 2023, approximately one-fifth of the population was affected by the XBB wave^7,8^. It is critical to assess whether the immunity in these subpopulations from XBB reinfection remains protective. For those uninfected by XBB, understanding how the efficacy of XBB-containing booster vaccines compares to natural immunity from breakthrough infection is vital, especially considering the emergence of new variants like JN.1 and BA.2.87.1.

## RESULTS

In this study, we collected blood samples from two groups of individuals who had previously experienced BA.5/BF.7 breakthrough infections following three doses of inactivated vaccines. One group (n=20) experienced XBB reinfection, while the other (n=11) received the WSK-V102C vaccine. Samples were collected about 1-month post-infection or vaccination (Supplementary Table 2). To evaluate their serum neutralization potency and breadth, we employed a panel of SARS-CoV-2 pseudoviruses (PsVs), including the wildtype (WT), B.1.617.2, BA.2, BA.5, XBB.1.5, XBB.1.16, EG.5.1, HK.3, BA.2.86, JN.1, and BA.2.87.1. Our previous study^4^ revealed that individuals with BA.5/BF.7 breakthrough infections primarily had high neutralizing titers against WT, moderately reduced (∼2-fold) against Delta and early Omicron variants, but significantly lower (∼10-fold) against XBB sub-lineages and BA.2.86. Interestingly, sera from XBB reinfection displayed a shifted neutralization pattern, showing the highest geometric mean titer (GMT) of 3,343 and 2,232 against BA.5 and BA.2, respectively, *i*.*e*., about 2-3-fold higher than that against WT. However, these sera showed remarkably decreased titers against XBB descendant subvariants (*e*.*g*., XBB.1.5, XBB.1.16, EG.5.1, HK.3) as well as BA.2.86 and its descendant JN.1, approximately 1.2-1.8-fold lower than that against WT. Notably, the lowest titer was observed against the latest variant, BA.2.87.1, with a GMT below 500 (Fig. 2a).

**Fig. 2.**
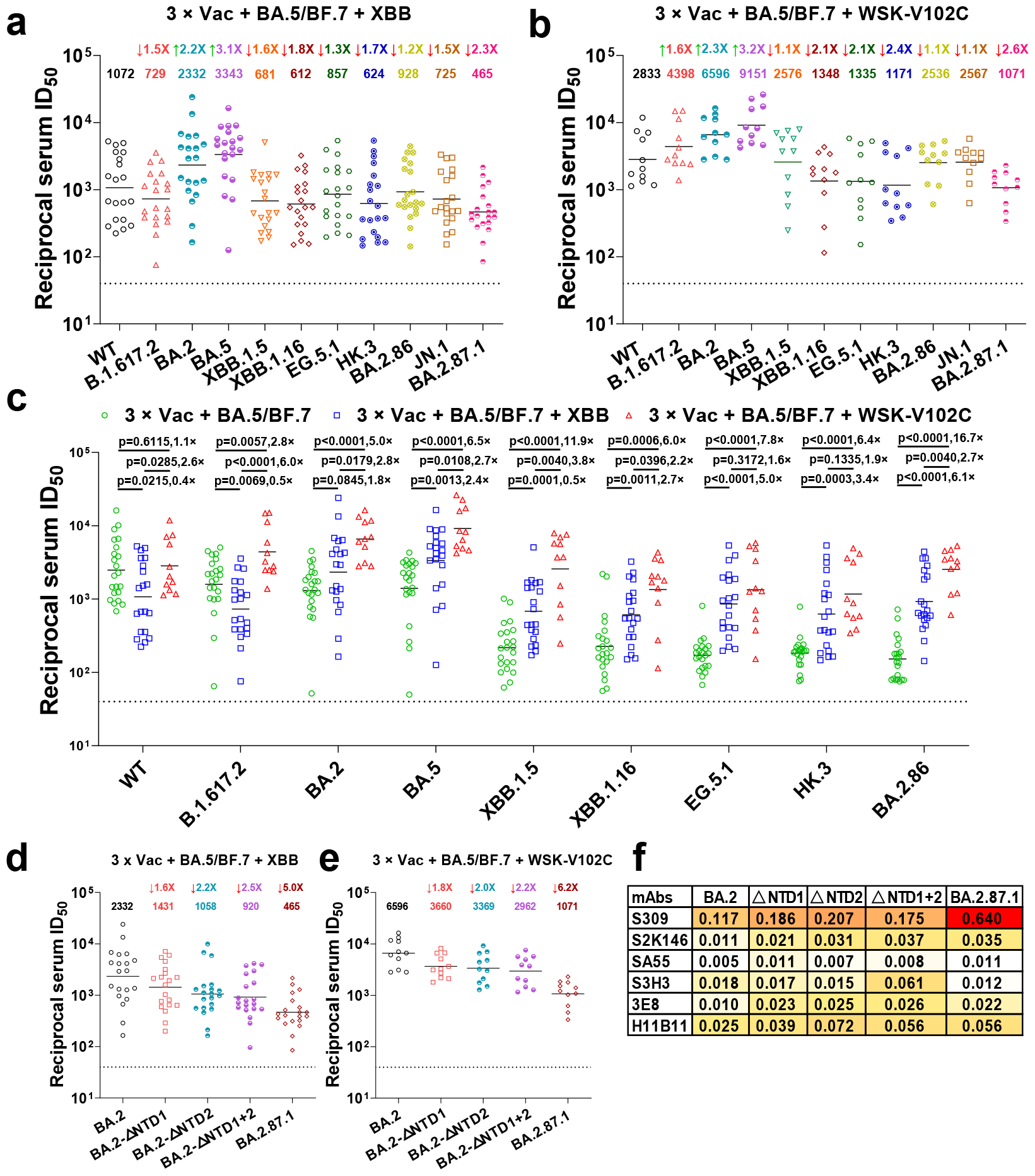
Neutralization of distinct SARS-CoV-2 sub-lineages by XBB reinfection and WSK-V102C vaccination sera. Neutralization of different SARS-CoV-2 variant PsVs by sera collected from two groups of individuals who had previously experienced BA.5/BF.7 breakthrough infections following three doses of inactivated vaccines: one group were reinfection with XBB virus (**a**), the other group were boosted with the WSK-V102C vaccine (**b**). **c** In parallel comparison of neutralization GMTs against distinct Omicron subvariants by sera collected from individuals with XBB reinfection, or with WSK-V102C booster vaccinations. Neutralization of PsVs representing BA.2.87.1’s single and combined NTD deletions by sera collected from individuals with XBB reinfection (**d**), or with WSK-V102C booster vaccination (**e**). **f** The neutralization IC50 (μg/ml) of mAbs against Omicron sub-lineage PsVs, presented as a heat map with darker colors implying greater change. Values above the symbols denote GMTs and their fold increase or decrease relative to WT (for panels A-B) or BA.2 (for panels D-E). Dotted lines indicate the threshold of detection (40 for all the cohorts). P values were determined by using Multiple Mann–Whitney tests. WT wild-type.

In contrast, sera from the group having received the WSK-V102C vaccine exhibited markedly higher neutralizing titers against all the tested variants. The highest neutralization titers were observed against BA.5 (GMT=9,151), followed by BA.2 (GMT=6,596) and Delta (GMT=4,398). While these sera also demonstrated reduced titers against XBB descendant subvariants, BA.2.86, JN.1, and BA.2.87.1, they maintained higher neutralization titers compared to the sera from the XBB reinfection group, although BA.2.87.1 exhibited a significantly higher escape potential than BA.2.86 and JN.1 (Supplementary Fig. 1). Even so, the sera from individuals vaccinated with the WSK-V102C vaccine maintained a GMT above 1,000 against BA.2.87.1 (Fig. 2b), suggesting that booster vaccination with WSK-V102C is expected to be effective in protecting individuals from BA.2.87.1 infection. Interestingly, we observed distinct antibody responses in the participants of the three groups: 1) those with BA.5/BF.7 breakthrough infections^4^, 2) those with additional XBB reinfection, and 3) those vaccinated with WSK-V102C post-breakthrough infection. The XBB reinfection group showed recalled antibody responses against post-Omicron variants, but less against pre-Omicron variants like WT and Delta. In contrast, the trivalent vaccine induced significantly more potent and broader neutralization activity (Fig. 2c). The elevated titers against Omicron variants may be attributed to the BA.5 and XBB.1.5 components in the vaccine, while the heightened response to Delta variants may be due to the vaccine’s Delta component.

For the BA.2.87.1 variant, the most notable mutations are likely the two fragment deletions in NTD. To assess their impact on neutralization sensitivity, we constructed PsVs with each of these fragment deletions, and a combination of both, based on the spike protein of BA.2 variant. The sera from individuals with XBB reinfection demonstrated a 1.6-fold reduction in neutralizing titer against the BA.2-ΔNTD1 (Del 15-26) PsV and a 2.2-fold reduction against BA.2-ΔNTD2 (Del 136-146) PsV, compared to the parental BA.2 PsV. Additionally, the PsV carrying spike protein with both fragment deletions (BA.2-ΔNTD1+2) showed an even further reduced sensitivity, at 2.5-fold lower than BA.2 PsV (Fig. 2d). A similar pattern was observed with sera from the individuals having received WSK-V102C vaccine (Fig. 2e), indicating that both fragment deletions in NTD contribute to the immune evasion of BA.2.87.1. However, as expected, these fragment deletions in NTD did not impact the efficacy of monoclonal antibodies (mAbs) targeting regions outside the NTD, as demonstrated by mAbs previously reported^5,9^ to retain activity against BA.2 (S309, S2K146, SA55, S3H3), nor did they affect ACE2-targeting mAbs, 3E8 and H11B11 (Fig. 2f).

## DISCUSSION

The rapid evolution of SARS-CoV-2 variants poses significant challenges in understanding and combating the virus. This study focuses on serum neutralization capacity against the newly emerged variants, particularly JN.1 and BA.2.87.1, which is crucial for evaluating the efficacy of existing vaccines. Our results show that sera from both XBB-reinfection and WSK-V102C-vaccination groups exhibited significantly lower neutralization titers against BA.2.87.1 than those against XBB sub-lineages, BA.2.86, and JN.1. We found that fragment deletions NTD in BA.2.87.1 spike protein made an important contribution to its escape from neutralization. However, BA.2.87.1 exhibited more significant resistance to neutralizing antibodies than the variant with combination of the two fragment deletions in NTD, suggesting that other mutations, like those in the RBD, also play a role in its immune evasion. This adds to our understanding of SARS-CoV-2’s immune evasion strategies.

The inclusion of participants with prior BA.5/BF.7 breakthrough infections and their responses to either XBB reinfection or the WSK-V102C vaccination provides valuable insights. Our study reveals that the WSK-V102C vaccine elicits more potent antibody responses compared to XBB reinfection, effectively against a broad spectrum of SARS-CoV-2 variants, which include Delta, BA.5, XBB sublineages, and notably, the more challenging subvariants JN.1 and BA.2.87.1. These findings suggest that all three components of the trivalent vaccine are functional and effective. Furthermore, the differential immune responses between individuals with XBB reinfection and those vaccinated with WSK-V102C highlight the complexity of immune memory and response to SARS-CoV-2, suggesting varied immunity patterns from natural infection and vaccination.

In summary, our study underscores the dynamic nature of SARS-CoV-2 evolution, particularly with the Omicron sub-lineages and emerging subvariants such as JN.1 and BA.2.87.1, which exhibit significant genetic divergence and enhanced escape capabilities. This situation calls for ongoing modifications in vaccine strategies. Updated vaccine formulations, such as SCTV01E we previously reported and WSK-V102C investigated in this study, demonstrate promising efficacy in generating high neutralizing titers against a spectrum of variants, including those with notable escape mutations. These findings highlight the crucial need for continuous monitoring and updating of vaccine formulations to keep pace with the rapidly evolving virus.

## DATA AVAILABILITY

All the data are provided in the main or the supplementary figures.

## ACKNOWLEDGMENTS

This study was supported by funding from the National Natural Science Foundation of China (32270142 to P.W.), Shanghai Rising-Star Program (22QA1408800 to P.W.), the Program of Science and Technology Cooperation with Hong Kong, Macao and Taiwan (23410760500 to P.W.), the National Key R&D Program of China (2023YFC2307800 to Shibo J.), Nanjing Research Center for Infectious Diseases of Integrated Traditional Chinese and Western Medicine (YBZX2022 to Y.Zhang), and NATCM’s Project of High-level Construction of Key TCM Disciplines. Pengfei Wang acknowledges support from AI for Science project of Fudan University (XM06231724), Open Research Fund of State Key Laboratory of Genetic Engineering, Fudan University (No. SKLGE-2304) and Xiaomi Young Talents Program.

## AUTHOR CONTRIBUTIONS

P.W., Y.Zhang, and M.L. designed and supervised the study; X.W., Shujun.J., and W.M. performed the experiments with help from C.Li and C.Liu; Shujun J., F.X., Y.Zhan, and J.Z. provided critical materials; P.W., Y.Zhang, M.L., X.W., Shujun J., Shibo J. and W.M. analyzed the data and wrote the manuscript. All authors reviewed, commented, and approved the manuscript.

## MATERIALS AND METHODS

### Serum samples

Blood samples from two groups of individuals who had previously experienced BA.5/BF.7 breakthrough infection following three doses of inactivated vaccines were collected at the Nanjing Hospital of Chinese Medicine. One group (n=20) experienced XBB reinfection, while the other (n=11) received the Trivalent XBB Vaccine (WSK-V102C). For all COVID-19 participants, the clinical diagnosis criteria were based on the ninth National COVID-19 guidelines. All participants involved in this study had mild symptoms. Their baseline characteristics are summarized in Supplementary Table 2. All the participants provided written informed consents. All collections were conducted according to the guidelines of the Declaration of Helsinki and approved by the ethical committee of Nanjing Hospital of Chinese Medicine Affiliated to Nanjing University of Chinese Medicine (number KY2023073).

### Cell lines

Expi293F cells (Thermo Fisher Cat# A14527) were cultured in the serum-free SMM 293-TI medium (Sino Biological Inc.) at 37 °C with 8% CO2 on an orbital shaker platform. HEK293T cells (Cat# CRL-3216), Vero E6 cells (cat# CRL-1586) were obtained from ATCC and cultured in 10% fetal bovine serum (FBS, GIBCO cat# 16140071) supplemented Dulbecco’s Modified Eagle Medium (DMEM, ATCC cat# 30-2002) at 37 °C, 5% CO2. I1 mouse hybridoma cells (ATCC, cat# CRL-2700) were cultured in Eagle’s Minimum Essential Medium (EMEM, ATCC cat# 30-2003) with 20% FBS.

### Monoclonal antibodies

Monoclonal antibodies tested in this study were constructed and produced in our laboratories at Fudan University. For each antibody, variable genes were optimized for human cell expression and synthesized by HuaGeneTM (China). VH and VL were inserted separately into plasmids (gWiz or pcDNA3.4) that encode the constant region for H chain and L chain. Monoclonal antibodies were expressed in Expi293F (ThermoFisher, A14527) by co-transfection of H chain and L chain expressing plasmids using Polyethylenimine and culture at 37 °C with shaking at 125 rpm and 8% CO2. On day 5, antibodies were purified using MabSelectTM PrismA (Cytiva, 17549801) affinity chromatography.

### Construction and production of variant pseudoviruses

Plasmids encoding the WT (D614G) SARS-CoV-2 spike and Omicron sub-lineage spikes, as well as the spikes with single or combined mutations were constructed. HEK293T cells were transfection with the indicated spike gene using Polyethylenimine (Polyscience). Cells were cultured overnight at 37°C with 5% CO2 and VSV-G pseudo-typed ΔG-luciferase (G*ΔG-luciferase, Kerafast) was used to infect the cells in DMEM at a multiplicity of infection of 5 for 4 h before washing the cells with 1×DPBS three times. The next day, the transfection supernatant was collected and clarified by centrifugation at 3000g for 10 min. Each viral stock was then incubated with 20% I1 hybridoma (anti-VSV-G; ATCC, CRL-2700) supernatant for 1 h at 37 °C to neutralize the contaminating VSV-G pseudotyped ΔG-luciferase virus before measuring titers and making aliquots to be stored at −80 °C.

### Pseudovirus neutralization assays

Neutralization assays were performed by incubating pseudoviruses with serial dilutions of monoclonal antibodies or sera, and scored by the reduction in luciferase gene expression. In brief, Vero E6 cells were seeded in a 96-well plate at a concentration of 2×10^4^ cells per well. Pseudoviruses were incubated the next day with serially diluted samples tested in triplicate for 30 min at 37 °C. The mixture was added to cultured cells and incubated for an additional 24 h. The luminescence was measured by Luciferase Assay System (Beyotime). IC50 was defined as the dilution at which the relative light units were reduced by 50% compared with the virus control wells (virus + cells) after subtraction of the background in the control groups with cells only. The IC50 values were calculated using nonlinear regression in GraphPad Prism.

### Sequence data collection

The sequencing data and corresponding genomic mutations of lineage BA.2.87.1 were retrieved from the GISAID database^10^ using the accession number provided in the Supplementary table 3. The global sequence metadata was retrieved from the CNGB RCoV19 database (ngdc.cncb.ac.cn/ncov/)^11^ on 2024 February 13 for the analysis of lineage frequency over time. The lineage mutation frequency data was retrieved from the outbreak.info website using its R package^12^.

### Construction of the phylogenetic tree

Sequences were sub-sampled from the Nextstrain GISAID global dataset of the last 6 months (nextstrain.org/ncov/gisaid/global/6m, updated on 2024 Feb 13) with approximately 8 to 20 sequences per clade. All 9 sequences from BA.2.87.1 were included in the analysis. The phylogenetic reconstruction was performed following the Nextstrain pipeline^13^. Visualization was done using iTOL^14^.

### Quantitative and statistical analysis

The statistical analyses for the pseudovirus neutralization assessments were performed using GraphPad Prism for calculation of mean value for each data point. Each specimen was tested in triplicate. Antibody neutralization IC50 values were calculated using a five-parameter dose-response curve in GraphPad Prism. For comparing the serum neutralization titers, statistical analysis was performed using Multiple Mann-Whitney tests. Two-tailed p values are reported. No statistical methods were used to determine whether the data met assumptions of the statistical approach.

