## Supplementary file for "Robust Neutralization of SARS-CoV-2 Variants Including JN.1 and BA.2.87.1 by Trivalent XBB Vaccine-Induced Antibodies"

**Supplementary Figure and Figure Legend**

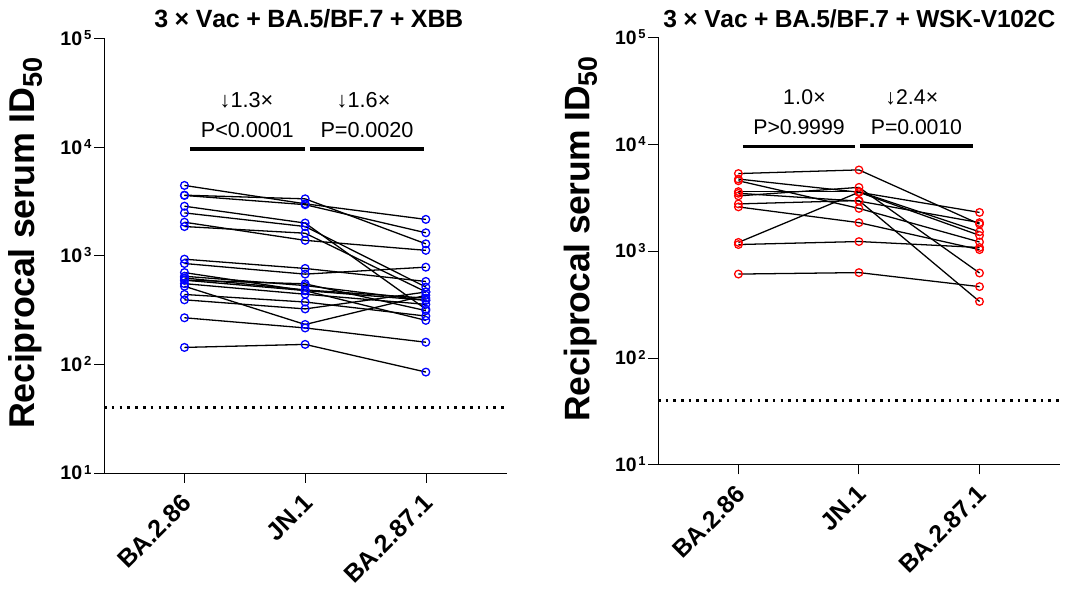
 **Supplementary Figure 1. In parallel comparison of neutralization titers from individuals with XBB reinfections and WSK-V102C booster vaccinations against BA.2.86, JN.1, and BA.2.87.1. p values were determined by using a Wilcoxon matched-pairs signed-rank test (two-tailed).**

**Supplementary Tables**

**Supplementary Table 1. The amino acid mutations of BA.2.87.1**

| **Mutation** | **Count** | **Gene** | **Ref** | **AApos** | **Var** |
| --- | --- | --- | --- | --- | --- |
| E T9I | 8 | E | T | 9 | I |
| E F23V | 9 | E | F | 23 | V |
| M D3N | 9 | M | D | 3 | N |
| M Q19E | 8 | M | Q | 19 | E |
| M A63T | 8 | M | A | 63 | T |
| M A104V | 8 | M | A | 104 | V |
| N P13L | 9 | N | P | 13 | L |
| N G30del | 1 | N | G | 30 | del |
| N E31del | 9 | N | E | 31 | del |
| N R32del | 9 | N | R | 32 | del |
| N S33D | 1 | N | S | 33 | D |
| N S33del | 8 | N | S | 33 | del |
| N Q43H | 1 | N | Q | 43 | H |
| N T148A | 9 | N | T | 148 | A |
| N G178D | 2 | N | G | 178 | D |
| N R203K | 9 | N | R | 203 | K |
| N G204R | 9 | N | G | 204 | R |
| N S413R | 8 | N | S | 413 | R |
| NS3 T32N | 9 | NS3 | T | 32 | N |
| NS3 T223I | 9 | NS3 | T | 223 | I |
| NS7a V71I | 1 | NS7a | V | 71 | I |
| NS8 A51V | 9 | NS8 | A | 51 | V |
| NS8 G66del | 9 | NS8 | G | 66 | del |
| NS8 S67del | 9 | NS8 | S | 67 | del |
| NS8 K68E | 9 | NS8 | K | 68 | E |
| NSP1 Q15R | 6 | NSP1 | Q | 15 | R |
| NSP1 M85del | 9 | NSP1 | M | 85 | del |
| NSP1 E93K | 9 | NSP1 | E | 93 | K |
| NSP1 S135R | 9 | NSP1 | S | 135 | R |
| NSP1 K141del | 1 | NSP1 | K | 141 | del |
| NSP1 S142del | 1 | NSP1 | S | 142 | del |
| NSP1 F143del | 1 | NSP1 | F | 143 | del |
| NSP12 E84K | 1 | NSP12 | E | 84 | K |
| NSP12 P323L | 9 | NSP12 | P | 323 | L |
| NSP12 G671S | 8 | NSP12 | G | 671 | S |
| NSP12 F694Y | 1 | NSP12 | F | 694 | Y |
| NSP12 L829F | 6 | NSP12 | L | 829 | F |
| NSP13 A134S | 2 | NSP13 | A | 134 | S |
| NSP13 V241L | 1 | NSP13 | V | 241 | L |
| NSP13 R392C | 9 | NSP13 | R | 392 | C |
| NSP13 Y421H | 1 | NSP13 | Y | 421 | H |
| NSP13 Q470K | 9 | NSP13 | Q | 470 | K |
| NSP13 A520V | 1 | NSP13 | A | 520 | V |
| NSP14 I42V | 5 | NSP14 | I | 42 | V |
| NSP14 L442S | 9 | NSP14 | L | 442 | S |
| NSP15 Q19H | 1 | NSP15 | Q | 19 | H |
| NSP15 T112I | 9 | NSP15 | T | 112 | I |
| NSP15 V148I | 1 | NSP15 | V | 148 | I |
| NSP2 T160I | 1 | NSP2 | T | 160 | I |
| NSP2 V594F | 1 | NSP2 | V | 594 | F |
| NSP3 T24I | 9 | NSP3 | T | 24 | I |
| NSP3 V170I | 1 | NSP3 | V | 170 | I |
| NSP3 I341T | 1 | NSP3 | I | 341 | T |
| NSP3 E392K | 1 | NSP3 | E | 392 | K |
| NSP3 G489S | 6 | NSP3 | G | 489 | S |
| NSP3 E545D | 9 | NSP3 | E | 545 | D |
| NSP3 S554A | 9 | NSP3 | S | 554 | A |
| NSP3 V766M | 2 | NSP3 | V | 766 | M |
| NSP3 P822L | 9 | NSP3 | P | 822 | L |
| NSP3 Y1463H | 1 | NSP3 | Y | 1463 | H |
| NSP3 A1526V | 9 | NSP3 | A | 1526 | V |
| NSP3 M1556I | 4 | NSP3 | M | 1556 | I |
| NSP3 S1682F | 9 | NSP3 | S | 1682 | F |
| NSP3 E1767D | 1 | NSP3 | E | 1767 | D |
| NSP3 P1921L | 1 | NSP3 | P | 1921 | L |
| NSP4 V99M | 2 | NSP4 | V | 99 | M |
| NSP4 L264F | 6 | NSP4 | L | 264 | F |
| NSP4 L323F | 1 | NSP4 | L | 323 | F |
| NSP4 T327I | 9 | NSP4 | T | 327 | I |
| NSP4 T492I | 9 | NSP4 | T | 492 | I |
| NSP4 T495A | 9 | NSP4 | T | 495 | A |
| NSP5 T24P | 1 | NSP5 | T | 24 | P |
| NSP5 P132H | 9 | NSP5 | P | 132 | H |
| NSP6 S106del | 9 | NSP6 | S | 106 | del |
| NSP6 G107del | 9 | NSP6 | G | 107 | del |
| NSP6 F108del | 9 | NSP6 | F | 108 | del |
| NSP7 L56F | 9 | NSP7 | L | 56 | F |
| NSP9 K36Q | 9 | NSP9 | K | 36 | Q |
| NSP9 D47N | 1 | NSP9 | D | 47 | N |
| NSP9 P71S | 9 | NSP9 | P | 71 | S |
| Spike C15del | 9 | Spike | C | 15 | del |
| Spike V16del | 9 | Spike | V | 16 | del |
| Spike N17del | 9 | Spike | N | 17 | del |
| Spike L18del | 9 | Spike | L | 18 | del |
| Spike T19del | 9 | Spike | T | 19 | del |
| Spike T20del | 9 | Spike | T | 20 | del |
| Spike R21del | 9 | Spike | R | 21 | del |
| Spike T22del | 9 | Spike | T | 22 | del |
| Spike Q23del | 9 | Spike | Q | 23 | del |
| Spike L24del | 9 | Spike | L | 24 | del |
| Spike P25del | 9 | Spike | P | 25 | del |
| Spike P26del | 9 | Spike | P | 26 | del |
| Spike A27S | 9 | Spike | A | 27 | S |
| Spike H69del | 9 | Spike | H | 69 | del |
| Spike V70del | 9 | Spike | V | 70 | del |
| Spike G75D | 9 | Spike | G | 75 | D |
| Spike S98F | 9 | Spike | S | 98 | F |
| Spike V126A | 9 | Spike | V | 126 | A |
| Spike C136del | 9 | Spike | C | 136 | del |
| Spike N137del | 9 | Spike | N | 137 | del |
| Spike D138del | 9 | Spike | D | 138 | del |
| Spike P139del | 9 | Spike | P | 139 | del |
| Spike F140del | 9 | Spike | F | 140 | del |
| Spike L141del | 9 | Spike | L | 141 | del |
| Spike G142del | 9 | Spike | G | 142 | del |
| Spike V143del | 9 | Spike | V | 143 | del |
| Spike Y144del | 9 | Spike | Y | 144 | del |
| Spike Y145del | 9 | Spike | Y | 145 | del |
| Spike H146del | 9 | Spike | H | 146 | del |
| Spike W152L | 9 | Spike | W | 152 | L |
| Spike F157L | 1 | Spike | F | 157 | L |
| Spike R190S | 9 | Spike | R | 190 | S |
| Spike V213G | 9 | Spike | V | 213 | G |
| Spike D215G | 9 | Spike | D | 215 | G |
| Spike G339D | 9 | Spike | G | 339 | D |
| Spike S371F | 9 | Spike | S | 371 | F |
| Spike S373P | 9 | Spike | S | 373 | P |
| Spike S375F | 9 | Spike | S | 375 | F |
| Spike T376A | 9 | Spike | T | 376 | A |
| Spike D405N | 9 | Spike | D | 405 | N |
| Spike R408S | 7 | Spike | R | 408 | S |
| Spike K417T | 8 | Spike | K | 417 | T |
| Spike N440K | 8 | Spike | N | 440 | K |
| Spike K444N | 8 | Spike | K | 444 | N |
| Spike V445G | 8 | Spike | V | 445 | G |
| Spike L452M | 8 | Spike | L | 452 | M |
| Spike N460K | 7 | Spike | N | 460 | K |
| Spike S477N | 9 | Spike | S | 477 | N |
| Spike N481K | 9 | Spike | N | 481 | K |
| Spike E484A | 9 | Spike | E | 484 | A |
| Spike Q498R | 9 | Spike | Q | 498 | R |
| Spike N501Y | 9 | Spike | N | 501 | Y |
| Spike Y505H | 9 | Spike | Y | 505 | H |
| Spike D614G | 9 | Spike | D | 614 | G |
| Spike P621S | 9 | Spike | P | 621 | S |
| Spike R634H | 1 | Spike | R | 634 | H |
| Spike V642G | 9 | Spike | V | 642 | G |
| Spike H655Y | 9 | Spike | H | 655 | Y |
| Spike N679R | 9 | Spike | N | 679 | R |
| Spike P681H | 9 | Spike | P | 681 | H |
| Spike S691P | 7 | Spike | S | 691 | P |
| Spike N764K | 6 | Spike | N | 764 | K |
| Spike T791I | 9 | Spike | T | 791 | I |
| Spike D796H | 9 | Spike | D | 796 | H |
| Spike D936G | 9 | Spike | D | 936 | G |
| Spike Q954H | 9 | Spike | Q | 954 | H |
| Spike N969K | 9 | Spike | N | 969 | K |

**Supplementary Table 2. Baseline characteristics of enrolled participants**

|  | WSK-V102C vaccine (n=11) | XBB reinfection individuals (n=20) |
| --- | --- | --- |
| Age(years), median(range) | 50.45(25-73) | 32(18-69) |
| Male, n(%) | 5(45.45%) | 7(35%) |
| BMI(kg/m^2^), mean(SD) | 23.11(3.48) | 22.54(4.34) |
| Breakthrough infections days after the last Coronavirus vaccines, median(range) | 397.45(358-420) | 400.07(113-707) |
| Number of Coronavirus vaccines doses | 4 | 3 |
| Days after the last Coronavirus vaccines or infection, median(range) | 39(27-58) | 19.8(7-36) |
| COVID-19 infection times | 1 | 2 |
| COVID-19 infection type | BA.5/BF.7 | BA.5/BF.7 or XBB |
| Comorbidities(%) |  |  |
| Any, n(%) | 0(0%) | 1(5%) |
| HTN, n(%) | 1(9.09%) | 3(15%) |
| CAD, n(%) | 0(0%) | 0(0%) |
| DM, n(%) | 0(0%) | 0(0%) |
| NAFLD, n(%) | 0(0%) | 0(0%) |
| Hyperlipidemia, n(%) | 0(0%) | 0(0%) |
| Obesity, n(%) | 2(18.18%) | 2(10%) |
| Arrhy, n(%) | 0(0%) | 0(0%) |
| Asthma, n(%) | 0(0%) | 0(0%) |
| Rhinitis, n(%) | 0(0%) | 0(0%) |
| Urticaria, n(%) | 0(0%) | 0(0%) |

BMI, body mass index. CAD, coronary artery disease. HTN, hypertension. DM, diabetes mellitus. Arrhy, arrhythmia. NAFLD, non-alcoholic fatty liver.

**Supplementary Table 3. GISAID accession number of 9 BA.2.87.1 sequences**

| **Strain** | **GISAID accession** | **Collection date** | **Country** |
| --- | --- | --- | --- |
| hCoV-19/SouthAfrica/NICD-R13515/2023 | EPI_ISL_18845398 | 2023/12/12 | South Africa |
| hCoV-19/SouthAfrica/NICD-R13200/2023 | EPI_ISL_18849984 | 2023/11/30 | South Africa |
| hCoV-19/SouthAfrica/NICD-N56614/2023 | EPI_ISL_18849985 | 2023/9/20 | South Africa |
| hCoV-19/SouthAfrica/NICD-N56836/2023 | EPI_ISL_18849986 | 2023/10/7 | South Africa |
| hCoV-19/SouthAfrica/NICD-N57176/2023 | EPI_ISL_18849987 | 2023/11/2 | South Africa |
| hCoV-19/SouthAfrica/NICD-N57208/2023 | EPI_ISL_18849988 | 2023/11/13 | South Africa |
| hCoV-19/SouthAfrica/NICD-N57216/2023 | EPI_ISL_18849989 | 2023/11/12 | South Africa |
| hCoV-19/SouthAfrica/NICD-N57440/2023 | EPI_ISL_18849990 | 2023/10/21 | South Africa |
| hCoV-19/SouthAfrica/NICD-N57469/2023 | EPI_ISL_18849991 | 2023/11/21 | South Africa |
